# PROTAC-mediated dual degradation of BCL-xL and BCL-2 is a highly effective therapeutic strategy in small-cell lung cancer

**DOI:** 10.1101/2024.02.27.582353

**Authors:** Sajid Khan, Lin Cao, Janet Wiegand, Peiyi Zhang, Maria Zajac-Kaye, Frederic J. Kaye, Guangrong Zheng, Daohong Zhou

## Abstract

BCL-xL and BCL-2 are validated therapeutic targets in small-cell lung cancer (SCLC). Targeting these proteins with navitoclax (formerly ABT263, a dual BCL-xL/2 inhibitor) induces dose-limiting thrombocytopenia through on-target BCL-xL inhibition in platelets. Therefore, platelet toxicity poses a barrier in advancing the clinical translation of navitoclax. We have developed a strategy to selectively target BCL-xL in tumors, while sparing platelets, by utilizing proteolysis-targeting chimeras (PROTACs) that hijack the cellular ubiquitin proteasome system for target ubiquitination and subsequent degradation. In our previous study, the first-in-class BCL-xL PROTAC, called DT2216, was shown to have synergistic antitumor activities when combined with venetoclax (formerly ABT199, BCL-2-selective inhibitor) in a BCL-xL/2 co-dependent SCLC cell line, NCI-H146 (hereafter referred to as H146), *in vitro* and in a xenograft model. Guided by these findings, we evaluated our newly developed BCL-xL/2 dual degrader, called 753b, in three BCL-xL/2 co-dependent SCLC cell lines and the H146 xenograft models. 753b was found to degrade both BCL-xL and BCL-2 in these cell lines. Importantly, it was considerably more potent than DT2216, navitoclax, or DT2216+venetoclax to reduce the viability of BCL-xL/2 co-dependent SCLC cell lines in cell culture. *In vivo*, 5 mg/kg weekly dosing of 753b leads to significant tumor growth delay similar to the DT2216+venetoclax combination in H146 xenografts by degrading both BCL-xL and BCL-2. Additionally, 753b administration at 5 mg/kg every four days induced tumor regressions. 753b at this dosage was well tolerated in mice without induction of severe thrombocytopenia as seen with navitoclax nor induced significant changes in mouse body weights. These results suggest that the BCL-xL/2 dual degrader could be an effective and safe therapeutic for a subset of SCLC patients warranting clinical trials in future.

## INTRODUCTION

Small-cell lung cancer (SCLC) is a highly aggressive malignancy with an overall 5-year survival rate of below 7%.(1) It remains a formidable clinical challenge secondary to a lack of effective targeted therapies and acquired resistance to conventional platinum-based chemotherapy i.e., the doublet of etoposide with cisplatin or carboplatin.(1–4) Despite advances in our understanding of the molecular landscape of SCLC, therapeutic breakthroughs have been limited. Recently, a PD-L1 inhibitor (atezolizumab) in combination with carboplatin and etoposide has been approved to treat extensive stage (advanced) SCLC.(5) However, this chemoimmunotherapy regimen has shown only a modest improvement in overall survival of patients compared to the chemotherapy alone.(3, 6) In June 2020, lurbinectedin, that works by inhibiting RNA polymerase II, has been approved by the FDA for the treatment of metastatic SCLC patients who show disease progression on or after platinum-based chemotherapy.(7, 8) However, lurbinectedin showed suboptimal responses in relapsed SCLC.(9) Thus, there is a critical unmet need for newer therapeutic strategies to effectively treat SCLC.

The BCL-2 family of proteins, including pro-apoptotic and anti-apoptotic (or pro-survival), are crucial regulators of intrinsic (mitochondrial) pathway of apoptosis.(10, 11) They have emerged as pivotal biomarkers in the survival of SCLC cells.(12–15) Among the anti-apoptotic members of the BCL-2 family, BCL-2, BCL-xL, and MCL-1, have garnered significant attention due to their prominent roles in promoting cell survival, chemoresistance, and disease progression in SCLC.(16–21) While individual targeting of these proteins has shown promise in preclinical studies, the development of resistance mechanisms and compensatory signaling pathways has hampered the success of monotherapies.(17, 20, 22–24) In recent years, a growing body of evidence has suggested that simultaneous inhibition of BCL-xL and BCL-2 or MCL-1 may represent a more effective strategy to combat SCLC.(12, 16, 18, 21, 25) Indeed, we have recently reported that the combined targeting of BCL-xL and MCL-1 with a platelet-sparing BCL-xL proteolysis targeting chimera (PROTAC) degrader (DT2216) and an mTOR inhibitor (AZD8055), is effective to inhibit tumor growth in SCLC preclinical models without causing on-target toxicities associated with BCL-xL and MCL-1 inhibitors.(26) In the same study, we found that a large subset of SCLC cell lines is co-dependent on BCL-xL and BCL-2 for their survival as evident from their high sensitivity to BCL-xL/2 inhibitor navitoclax (formerly ABT263). Unfortunately, navitoclax causes dose-limiting thrombocytopenia through on-target inhibition of BCL-xL in platelets. The severe platelet toxicity presents a great challenge in the clinical translation of navitoclax and other BCL-xL inhibitors.(27–29) Therefore, developing strategies to reduce BCL-xL inhibition-induced platelet toxicity has been an important research area for more than a decade now.(30) One of the earliest approaches to reduce BCL-xL inhibition-induced platelet toxicity was to convert a potent BCL-xL inhibitor into a pro-drug such as APG-1252.(31) The idea behind a pro-drug of APG-1252 is to minimize its impact on platelets in the bloodstream until it reaches the tumor site, where it is then converted into its active form. Alternatively, we have used the PROTAC technology to reduce navitoclax thrombocytopenia by converting it into a VHL E3 ligase targeted BCL-xL PROTAC, such as DT2216, because platelets are devoid of VHL E3 ligase required for BCL-xL degradation.(32) In contrast, these PROTACs can efficiently degrade BCL-xL in tumor cells, which possess significantly high levels of VHL compared to platelets.(32–35)

Through extensive structural optimizations, we have developed new PROTACs that are capable of degrading both the BCL-xL and BCL-2.(36) The lead first-in-class BCL-xL and BCL-2 dual degrader, named 753b, was shown to more potently degrade BCL-xL compared to DT2216 with concomitant degradation of BCL-2. This was due to accessibility of a key lysine residue in BCL-2 that was not accessible by DT2216 as well the formation of a more stable ternary complex between VHL E3 ligase, 753b and BCL-xL or BCL-2. By degrading both the proteins, 753b was found to be more potent in killing of BCL-xL/2 co-dependent cancer cells.(36)

In this study, we capitalize on the synergistic effects of degrading BCL-xL and BCL-2 in SCLC using a single degrader i.e., 753b.(36) Since a large subset of SCLC are characterized by high BCL-xL and BCL-2 mRNA and protein expression, we evaluated 753b in comparison to DT2216, navitoclax and DT2216+venetoclax combination on efficiency and specificity for BCL-xL and BCL-2 degradation as well as viability of SCLC cells. Next, we evaluated anti-tumor efficacy of 753b in NCI-H146 (hereafter referred to as H146) SCLC xenograft models at two different dosing frequencies and tumor sizes. The degradations of BCL-xL and BCL-2 following treatment with 753b were confirmed in tumor tissues. Collectively, our results show that 753b is significantly more potent than DT2216, navitoclax and the DT2216+venetoclax combination in killing BCL-xL/2 co-dependent SCLC cells by degradation of both the proteins. *In vivo*, 753b requires significantly lower dosage to elicit similar effects as DT2216+venetoclax and is capable of regressing larger H146 xenograft tumors.

## MATERIALS AND METHODS

### Cell lines and culture

All the SCLC cell lines, except H146 (hereafter all the SCLC cell lines in this article are referred to without the prefix ‘NCI’), were obtained from the original NCI-Navy Medical Oncology source supply.(37) H146 cells were obtained from the American Type Culture Collections (ATCC, Manassas, VA). SCLC cell lines were cultured in RPMI-1640 medium (Cat. No. 22400–089, Thermo Fisher, Waltham, MA). The culture media were supplemented with 10% heat-inactivated fetal bovine serum (FBS, Cat. No. S11150H, Atlanta Biologicals, GA), 100 U/mL penicillin and 100 µg/mL streptomycin (Pen-Strep, Cat. No. 15140122, Thermo Fisher). The stocks of NCI SCLC cell lines that we used were STR profiled by NIH or the ATCC, so the authenticity of the cell lines remained preserved. WI38 normal lung fibroblasts were obtained from ATCC and were cultured in high-glucose DMEM medium (Cat. No. 12430062, Thermo Fisher), supplemented with FBS and pen-strep. All cultures were confirmed for Mycoplasma negativity using the MycoAlert Mycoplasma Detection Kit (Cat. No. LT07– 318). All the cell lines were maintained in a humidified incubator at 37 °C and 5% CO_2_.

### Chemical compounds

DT2216 and 753b were synthesized in Dr. Guangrong Zheng’s laboratory (University of Florida, Gainesville, FL) according to the previously described protocol.(32, 36) Navitoclax (Cat. No. S1001) and venetoclax (Cat. No. S8048) were purchased from SelleckChem (Houston, TX). All the compounds were dissolved in DMSO at 10 mM stock solution for *in vitro* assays. The details of *in vivo* formulations are provided in the ‘Tumor xenograft studies’ method.

### Cell viability assays

Cells were seeded in 96-well plates at a density of 5×10^3^ (adherent cells) or 5×10^4^ (suspension cells) per well, and then treated the following day (adherent cells) or the same day (suspension cells) in 9-point 3-fold serial dilutions of the drugs in 3 or 6 replicates. The cell viability was measured by adding MTS reagent (Cat. No. G-111, Promega, Madison, WI) according to the manufacturer’s protocol and as described previously.(26, 32) The absorbance was recorded at 490 nm using Biotek’s Synergy Neo2 multimode plate reader (Biotek, Winooski, VT). The half-maximal inhibitory concentration (IC_50_) values were determined using GraphPad Prism software (GraphPad Software, La Jolla, CA).

### Immunoblotting

Cells or tumor tissues were lysed using RIPA lysis buffer (Cat. No. BP-115DG, Boston Bio Products, Ashland, MA) supplemented with protease and phosphatase inhibitor cocktail (Cat. No. PPC1010, Sigma-Aldrich, St. Louis, MO) as described previously.(26, 32) Briefly, an equal amount of protein samples (20-40 µg/lane) were loaded to precast gels and transferred onto PVDF membranes. The membranes were blocked with 5% (w/v) non-fat dry milk in TBST buffer, and subsequently probed with primary antibodies overnight at 4 °C. After washing with TBST, the membranes were incubated with horseradish peroxidase (HRP)-linked secondary antibody for 1-2 h at room temperature. Finally, the membranes were incubated with chemiluminescent HRP substrate (Cat. No. WBKLS0500, Millipore Sigma, Billerica, MA), and were recorded using the ChemiDoc MP Imaging System (Bio-Rad, Hercules, CA). The immunoblots were quantified using Image J software. The primary antibody details are provided in *Supplementary Table 1*.

### Tumor xenograft studies

CB-17 SCID-beige mice aged 5-6 weeks were purchased from the Charles River Laboratories (Wilmington, MA). H146 cells at a concentration of 5×10^6^ per mouse in 50% Matrigel (Cat. No. 356237, Corning, Corning, NY) mixed with 50% serum/antibiotic-free RPMI medium were injected subcutaneously (s.c.) into the right flank region of the mice as described previously.(26, 32) Tumor size was measured twice a week with digital calipers and tumor volume was calculated using the formula (Length×Width^2^×0.5). The mice were randomized into different treatment groups when the tumors reached ∼150 mm^3^ (for tumor inhibition study) or ∼500 mm^3^ (for tumor regression study). Mice were treated with vehicle, 753b (5 mg/kg, once a week or every four days, i.p.), or DT2216 (15 mg/kg, once a week, i.p.). DT2216 and 753b were formulated in 50% phosal 50 PG, 45% miglyol 810N and 5% polysorbate 80. Post-euthanasia, the tumors were harvested, lysed, and used for immunoblotting analysis. All the animal experiments were performed in accordance with the IACUC policies.

### Platelet counts

Approximately 50 µL of blood was collected from each mouse in EDTA-treated tubes *via* the *submandibular plexus*. The blood was immediately used for platelet counts using an automated hematology analyzer HEMAVET 950FS (Drew Scientific Inc., Miami Lakes, FL). The data were expressed as number of platelets per µL of blood.

### Statistical Analysis

For analysis of the means of three or more groups, analysis of variance (ANOVA) test was performed. In the event that ANOVA justified post-hoc comparisons between group means, the comparisons were conducted using Tukey’s multiple-comparisons test. A two-sided unpaired Student’s *t*-test was used for comparisons between the means of two groups. *P*<0.05 was considered to be statistically significant.

## RESULTS

### 753b is a dual BCL-xL/2 degrader in SCLC cells

In our previous study, we conducted a BH3 mimetic screening, where SCLC cell lines were treated with selective inhibitors of BCL-2, BCL-xL and MCL-1 and a BCL-xL/2 dual inhibitor to determine their survival dependencies. Through this screening, we determined that SCLC cell lines are heterogenous in their dependence on BCL-2 family proteins, and we defined a subset of SCLC tumors that depended on both BCL-2 and BCL-xL for survival.(26)

The BCL-xL/2 dual inhibitor navitoclax is highly effective in SCLC as evidenced by CancerRxGene data, where about 70% of the cell lines were found to be sensitive.(38) Similar results were obtained by BH3 mimetic screening performed by us in a previous study.(26) In that screening, we also found that about 45% of the tested SCLC cell lines are sensitive to BCL-xL-selective inhibitor A1155463 and about 70% are sensitive to navitoclax. These observations suggest that dual targeting of both BCL-xL and BCL-2 is a preferred therapeutic strategy in SCLC. Since navitoclax causes dose-limiting and on-target thrombocytopenia through inhibition of BCL-xL in platelets, it has not been further developed for clinical translation. To circumvent these challenges, we have recently identified a BCL-xL/2 dual degrader, named 753b, through extensive structural optimization of DT2216 (**Fig. 1a**).(36) However, the dual BCL-xL/2 degradation by 753b was not evaluated in SCLC cells in our previous proof-of-concept study. 753b required shorter exposure compared to DT2216 to induce BCL-xL degradation in SCLC cells, which typically grow in multicellular clusters (**Fig. 1b-d; Supplementary Fig. 1a & b**). Moreover, 753b exhibited higher cell line average D_max_ (% maximum degradation) and reduced average DC_50_ (concentration at which 50% degradation occurs) for BCL-xL in H146 and H211 cells compared to DT2216 (**Fig. 1e-g; Supplementary Fig. 1c & d**). Additionally, 753b induced substantial degradation of BCL-2 (D_max_: 26.3 to 62.7%) in SCLC cells (**Fig. 1b-g**). Among the SCLC cell lines tested, H211 cells showed maximum BCL-2 degradation followed by H1059 and H146, respectively. Notably, we observed a moderate MCL-1 suppression at higher doses of 753b in H146 cells, likely due to MCL-1 cleavage by activated caspase-3 as reported previously.(39–41) 753b treatment resulted in no considerable degradation of BCL-w. These results suggest that 753b is an effective dual degrader of BCL-xL and BCL-2 in SCLC cells.

**Figure 1.**
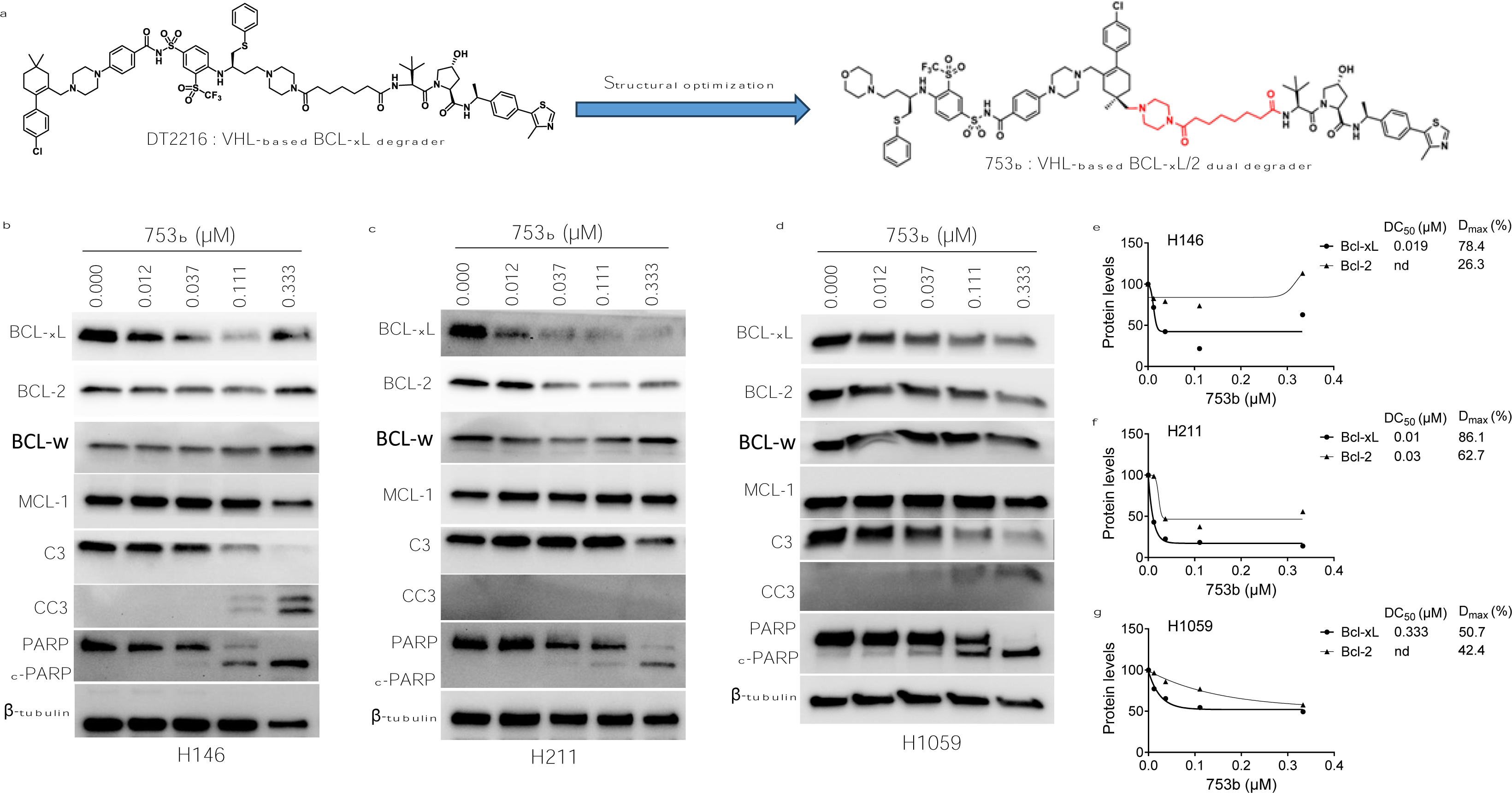
753b degrades both BCL-xL and BCL-2 leading to apoptosis in SCLC cells. **a,** Chemical structures of DT2216 and 753b. **b-d**, Immunoblot analyses of BCL-X_L_, BCL-2, BCL-w, MCL-1, caspase 3 (C3), cleaved caspase 3 (CC3), PARP and cleaved (c)-PARP in SCLC H146 (b), H211 (c) and H1059 cells (d) after they were treated with indicated concentrations of 753b for 24 h. The β-tubulin was used as an equal loading control. **e-g**, Densitometric analysis of BCL-xL and BCL-2 immunoblots in H146 (e), H211 (f) and H1059 cells (g) showing DC_50_ and D_max_ values for each protein. DC_50,_ concentration required to degrade 50% of the protein; D_max,_ maximum degradation in percentage; nd, not determined.

### 753b is highly potent to kill BCL-xL/2-dependent SCLC cells

The dual BCL-xL/2 degradation by 753b induced robust apoptosis as indicated by increased caspase-3 and PARP cleavage in SCLC cells (**Fig. 1b-d**). 753b exerted 12-, 15- and 5-fold higher potency than DT2216 to kill BCL-xL/2-dependent H146, H211 and H1059 cells, respectively. Moreover, it was 2- and 4-fold more potent than navitoclax against H146 and H211 cells, respectively (**Fig. 2a-c & f**). We also tested the effect of 753b on the viability of BCL-xL/MCL-1-dependent H378 cells and WI38 normal lung fibroblasts, which were found to be mildly sensitive and resistant to 753b, respectively (**Fig. 2d-f**). This suggests that 753b exerts BCL-xL/2-specific activity and possesses a wide therapeutic window.

**Figure 2.**
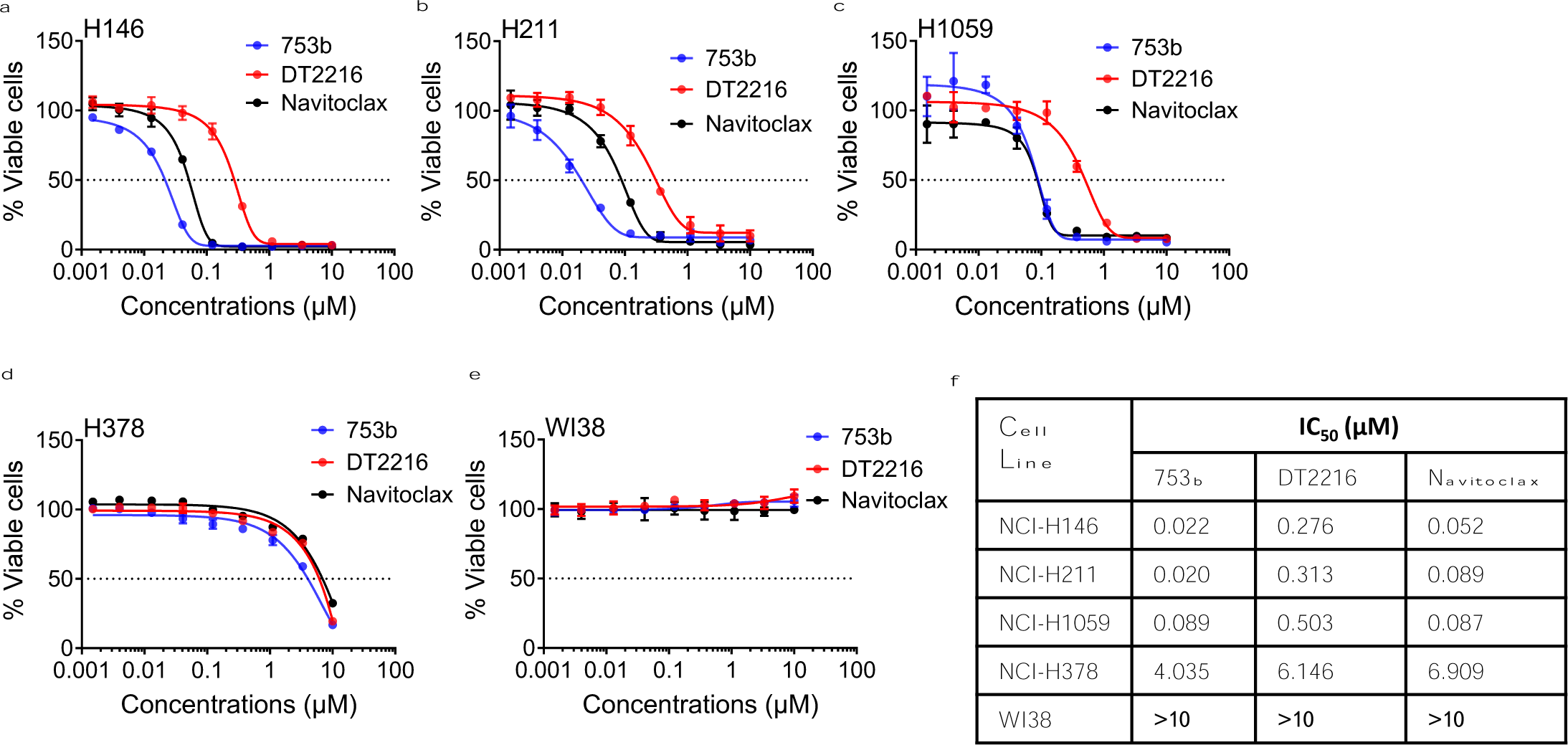
753b is more potent than DT2216 or navitoclax to kill BCL-X_L_/2-dependent SCLC cells. **a-e**, Viability of H146 (a), H211 (b), H1059 (c), H378 (d) SCLC cells and WI38 normal lung fibroblasts (e) after they were treated with increasing concentrations of 753b, DT2216 or navitoclax for 72 h. **f**, IC_50_ values for 753b, DT2216 and navitoclax in SCLC cell lines and WI38 cells are tabulated.

H146, H211 and H1059 cells were more sensitive to a combination of DT2216 with venetoclax compared to mono-targeting, confirming their co-dependence on BCL-xL and BCL-2 (**Fig. 3a-c**). Therefore, we tested whether venetoclax can similarly enhance the efficacy of 753b. We found that venetoclax could only slightly enhance the efficacy of 753b (**Fig. 3d-f**). This was expected because 753b cannot completely degrade BCL-2, so the slightly enhanced efficacy might have resulted from the inhibition of remaining BCL-2. However, 753b alone was equally effective as the combination of DT2216 and venetoclax against these SCLC cell lines. These results confirm that 753b is a dual BCL-xL/2 degrader that can be more potent against BCL-xL/2 co-dependent SCLC cells than DT2216 alone and does not require its combination with venetoclax to effectively kill these cells.

**Figure 3.**
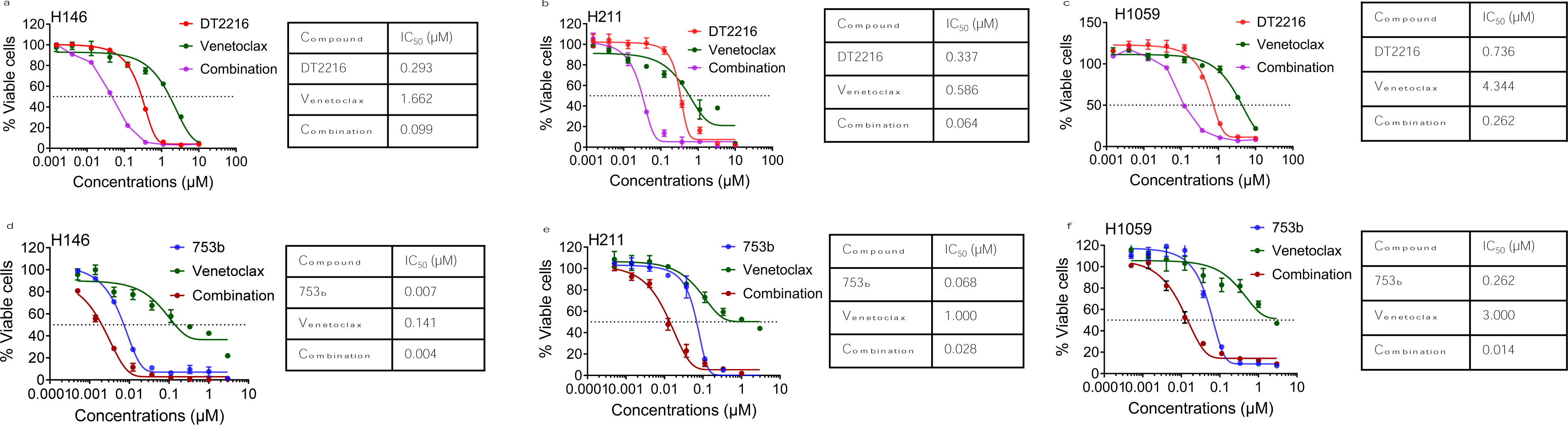
753b showed comparable or enhanced efficacy as DT2216+venetoclax combination to kill BCL-X_L_/2-dependent SCLC cells. **a-c,** Viability of H146 (a), H211 (b) and H1059 cells (c) after they were treated with increasing concentrations of DT2216, venetoclax or their 1:1 combination. IC_50_ values for the individual agents and combinations are tabulated. **d-f**, Viability of H146 (d), H211 (e) and H1059 cells (f) after they were treated with increasing concentrations of 753b, venetoclax or their 1:1 combination. IC_50_ values for the individual agents and combinations are tabulated.

### 753b showed similar anti-tumor efficacy as DT2216+venetoclax by dual BCL-xL/2 degradation in H146 xenograft model

Finally, we tested the *in vivo* efficacy and safety of 753b using H146 xenograft models. In the first study, tumor bearing mice were treated with 5 mg/kg weekly dosing of 753b compared with DT2216 at 15 mg/kg weekly alone or in combination with venetoclax (50 mg/kg, 5 days a week). Despite a 3-fold lower dosing, 753b led to similar tumor growth inhibition as DT2216 initially for five weeks, and thereon tumors grew slower in 753b-treated mice compared to DT2216-treated mice. Tumor growth after five weeks of treatment was similar in 753b- and DT2216+venetoclax-treated mice (**Fig. 4a**). In addition, mice treated with 753b showed significantly enhanced survival compared to vehicle control (median survival: 104 vs 32 days) and showed a trend towards extended survival compared to DT2216 (median survival: 104 vs 94 days) (**Fig. 4b**). The median survival in both 753b- and DT2216+venetoclax-treated groups was 104 days. None of these treatments caused any significant changes in mouse body weights (**Fig. 4c**). We also measured platelet levels one day after the first dose and three days after the third weekly dose. We observed that the platelet reduction one day after the first dose was more pronounced in 753b-treated mice than DT2216-treated mice, but similar reductions were observed three days after third dose. Addition of venetoclax to DT2216 caused no further reduction in platelets. Notably, these reductions in platelets did not reach clinically dangerous levels, i.e., below 1×10^5^ per µL of blood, and thus were well tolerable in mice (**Supplementary Fig. 2a & b**). These results suggest that 753b is a more potent antitumor agent than DT2216 and shows similar efficacy as the combination of DT2216+venetoclax against BCL-xL/2 co-dependent SCLC.

**Figure 4.**
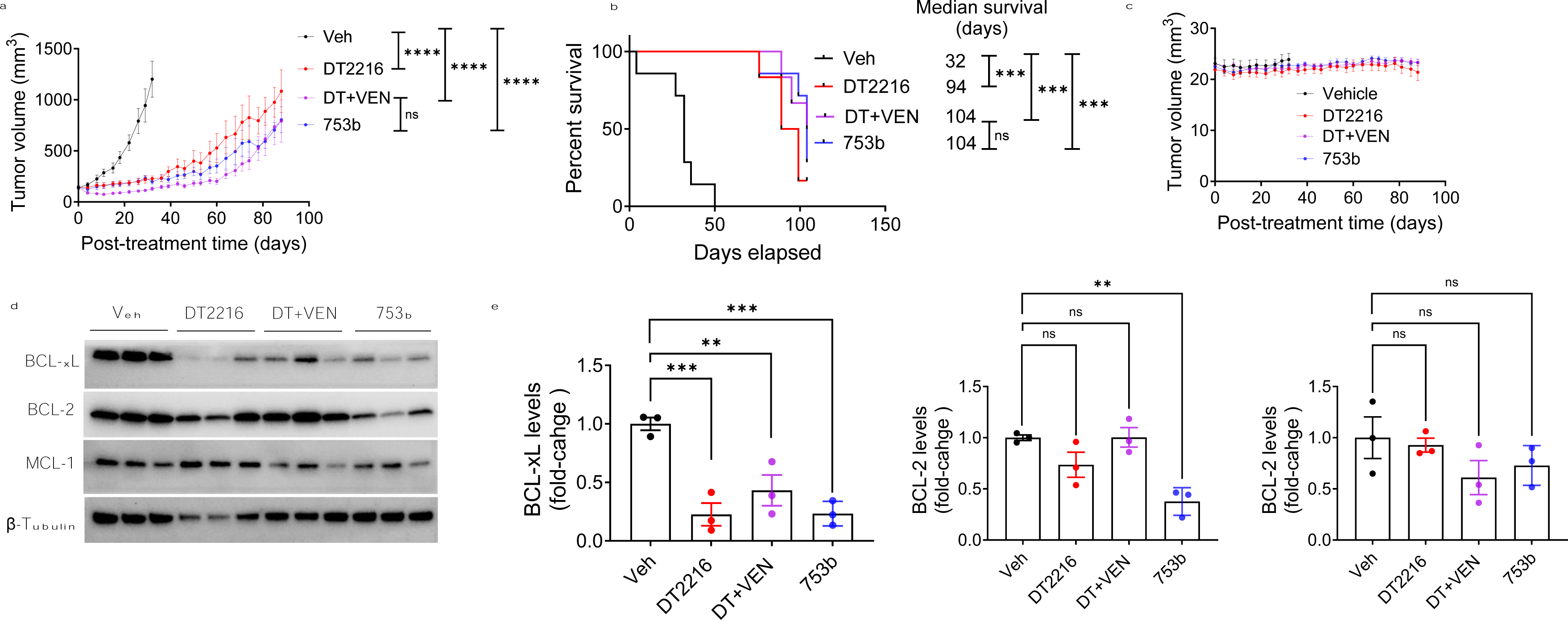
753b is more potent than DT2216 and similar to DT2216+venetoclax to inhibit growth of BCL-X_L_/2-dependent H146 xenograft tumors in mice. **a**, Tumor volume changes in H146 xenografts after treatment with vehicle, DT2216 (15 mg/kg, weekly i.e., q7d, i.p.), a combination of DT2216 with venetoclax (50 mg/kg, 5 days a week, p.o.), or 753b (5 mg/kg, q7d, i.p.). Data are presented as mean ± SEM (n = 7, 6, 6 and 7 mice in vehicle, DT2216, and DT2216+venetoclax, and 753b groups, respectively, at the start of treatment). When the biggest tumor dimension reached 1.5 cm, the mice were sacrificed in accordance with IACUC protocol, and the remaining mice were continued treated up to104 days. Tumor volume changes are shown up to post-treatment day 88, when 5 or more mice were alive in each treatment group. \*\*\*\**p* <0.0001, ns: not significant as determined by one-way ANOVA and Dunnett’s multiple comparisons test at post-treatment day 32. **b**, Kaplan-Meier survival analysis of mice as treated in **a**. Survival times was recorded at the tumor endpoint i.e., biggest tumor dimension of 1.5 cm or more. The median survival time is shown on the right. \*\*\**p* <0.001, ns: not significant as determined by two-sided Student’s t-test. **c**, Mouse body weight changes in H146 xenografts after treatment as in **a**. Data are presented as mean ± SEM. **d**, Immunoblot analysis of BCL-xL, BCL-2, and MCL-1 in H146 xenograft tumors two days after last treatment with vehicle, DT2216, DT2216+venetoclax (DT+VEN), or 753b (*n* = 3 mice per group) as in **a**. **e**, Densitometric analysis of immunoblots in **d**. \**p* <0.05, \*\**p* <0.01, \*\*\**p* <0.001 compared to vehicle as determined by one-way ANOVA and Dunnett’s multiple comparisons test.

We wondered whether 753b can induce degradation of both BCL-xL and BCL-2 in tumors as observed in *in vitro*. Indeed, 753b showed significant degradation of both BCL-xL and BCL-2 in excised xenograft tumors. As expected, DT2216 selectively degraded BCL-xL, without significant effect on BCL-2. Neither 753b, nor DT2216 showed any significant effect on MCL-1. These data suggest that the enhanced anti-tumor efficacy of 753b compared to DT2216 is derived from dual degradation of BCL-xL and BCL-2 in tumors (**Fig. 4d & e**).

### 753b treatment induced tumor regressions in H146 xenograft model

We also investigated whether 753b can regress larger established tumors in mice. For this, mice were treated every four days with 5 mg/kg dose of 753b when the average tumor sizes were more than 500 mm^3^. Interestingly, 753b caused significant tumor regressions in these mice. These regressions sustained for about three weeks and then tumors started to slowly regrow (**Fig. 5a**). The excised tumors were significantly smaller in 753b-treated mice compared to vehicle-treated mice (**Fig. 5b & c**). At this dosage, 753b caused no significant changes in mouse body weights, suggesting this dosing regimen is tolerated in mice (**Fig. 5d**).

**Figure 5.**
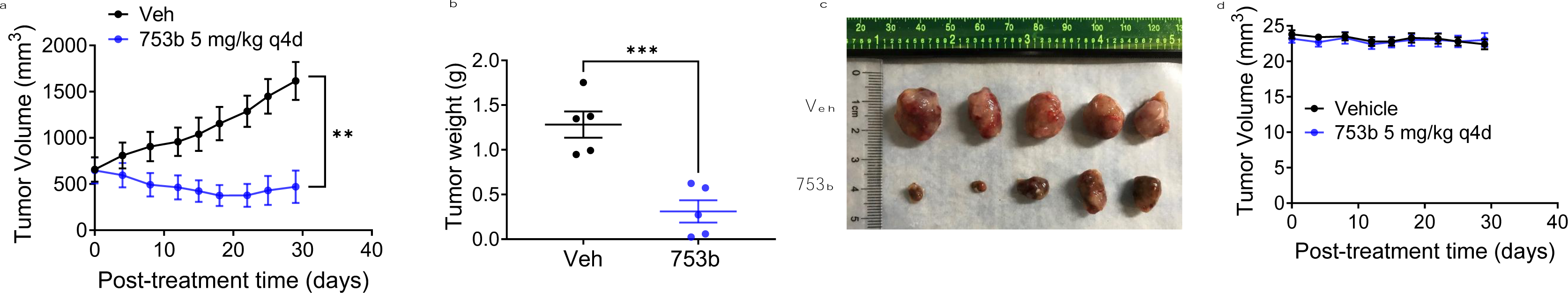
753b induces regressions of larger H146 xenograft tumors in mice. **a,** Tumor volume changes in H146 xenografts after treatment with vehicle, or 753b (5 mg/kg, every four days i.e., q4d, i.p.). Data are presented as mean ± SEM (n = 5 mice). \*\**p* <0.01 compared to vehicle as determined by two-sided Student’s t-test. **b**, Tumor weights at the end of experiment in **a**. Data are presented as mean ± SEM (n = 5 mice). \*\*\**p* <0.001 compared to vehicle as determined by two-sided Student’s t-test**. c**, The images of excised tumors from **a**. **d**, Mouse body weight changes in H146 xenografts after treatment as in **a.**

## DISCUSSION

The BCL-2 family of anti-apoptotic proteins, such as BCL-xL, BCL-2, and MCL-1, represent promising therapeutic targets for SCLC, a malignancy characterized by its aggressive nature and resistance to standard treatment modalities.(1–4, 12–15) Therefore, the targeting of BCL-xL and BCL-2 with small molecule inhibitors has been explored as a therapeutic strategy in SCLC, since the discovery of navitoclax.(1, 16–21) However, the only FDA approved BCL-2 family protein inhibitor for the treatment of relapsed/refractory chronic lymphoblastic leukemia (CLL) and acute myelogenous leukemia (AML) is an alternate BCL-2-selective agent, venetoclax.(42, 43) The single agent efficacy of venetoclax is limited in SCLC because only a very small subset of SCLC exhibit dependence on BCL-2 expression for survival, while a large subset of SCLC samples exhibit co-dependence on BCL-xL and BCL-2 or MCL-1.(12, 13, 16) Therefore, a more promising strategy to co-target BCL-xL and MCL-1 with DT2216 and AZD8055, respectively, in SCLC has been recently reported by us.(26) In the current study, we focus on co-targeting BCL-xL and BCL-2. Since the clinical drug development of the BCL-xL/2 dual inhibitor navitoclax has been hindered by dose-limiting and on-target thrombocytopenia due to BCL-xL inhibition in platelets,(27–29) we have recently converted navitoclax into platelet-sparing BCL-xL PROTAC degraders using VHL E3 ligase for their activity which significantly reduced platelet toxicity.(32) This is because VHL is minimally expressed in platelets, limiting the effect on platelets and leading to selective killing of tumor cells. (32–35)

Given the high potency of navitoclax, we proposed that a dual BCL-xL and BCL-2 degrader could have similar or improved efficacy with reduced platelet toxicity in SCLC compared to navitoclax. Our group has recently reported a BCL-xL/2 dual degrader, named 753b, that shows improved antileukemic activity compared to BCL-xL-selective degraders such as DT2216.(36) Therefore, we utilized 753b to test our hypothesis in SCLC. 753b was found to be 5- to 15-fold more potent than DT2216 in killing BCL-xL/2 co-dependent SCLC cells. Moreover, it was 2- to 4-fold more potent than navitoclax likely due to the catalytic mechanism of PROTACs. Furthermore, 753b induced deeper and rapid degradation of BCL-xL at significantly lower concentrations than DT2216, and concomitantly leads to BCL-2 degradation. However, the extent of BCL-2 degradation by 753b was cell line-dependent and was seen at slightly higher concentrations compared to BCL-xL degradation. Furthermore, we elucidated the downstream signaling events triggered by the dual degradation of BCL-xL and BCL-2. 753b leads to a strong activation of caspase 3 and PARP demonstrating a pronounced shift towards apoptotic pathways in SCLC cells. The high cell death at supra-IC_50_ concentrations of 753b leads to MCL-1 suppression possibly through a caspase 3-mediated cleavage. This is because MCL-1 is one of the substrates of caspase 3.(39–41) The observed increase in caspase activation and cleavage of key apoptotic substrates reinforces the notion that co-targeting BCL-xL and BCL-2 for degradation induces a robust pro-apoptotic response in SCLC cells, ultimately leading to reduced cell viability.

Further evaluations in H146 xenograft models demonstrate that 753b requires threefold lower dosage (5 mg/kg once a week) to elicit similar antitumor activity as the combination of DT2216+venetoclax. However, the tumors started slowly regrowing after five weeks of treatment, though the regrowth was slower in 753b- and DT2216+venetoclax treated mice compared to DT2216-treated mice. Also, 753b at this dosage leads to trend for extending median survival compared to DT2216, however the difference did not reach statistical significance. 753b was tolerated in mice with no measurable toxicity including no significant body weight loss nor induce severe platelet toxicity. The platelet reductions after 753b treatment never reached below 1×10^5^ per µL of blood, which is considered to be safe.(26, 32) Despite the lower dosing, 753b exerted significant BCL-xL and BCL-2 degradation in tumors collected at the end of treatment, which further supports the data showing comparable antitumor activity of 753b to the DT2216+venetoclax combination. Treatment with 753b and DT2216+venetoclax also resulted in moderate, but non-significant, reduction in MCL-1 levels. These results also suggest that the tumor regrowth in 753b- and DT2216-treated mice was not due to their inability to degrade BCL-xL and/or BCL-2 or compensatory elevation of MCL-1. Therefore, it would be crucial to elucidate the mechanisms responsible for tumor regrowth after a period of tumor statis upon 753b- and DT2216 treatment for designing combination strategies to treat SCLC more effectively in future studies.

Though it is a common laboratory practice to treat mice when their tumors are small in size (100-200 mm^3^), we are aware that the tumor sizes in human patients are larger at the initiation of therapy. Therefore, we tested whether 753b can inhibit or regress larger tumors in mice in a separate experiment using the H146 xenograft model. We initiated 753b treatment when the tumors were > 500 mm^3^. Since the tumors were significantly larger, we treated mice with 5 mg/kg of 753b every four days. 753b at this dosage was found to be efficient in regressing these larger H146 xenograft tumors. The tumor regressions sustained for three weeks and then started slowly growing, leading to a significant tumor growth delay without causing any changes in mouse body weights.

Our findings further support the hypothesis that dual targeting of BCL-xL and BCL-2 results in a synergistic pro-apoptotic effect, effectively disrupting the survival machinery of SCLC cells. This synergism is consistent with previous studies highlighting the compensatory mechanisms and functional redundancy within the BCL-2 family proteins, which may explain the limited success of monotherapies targeting either BCL-xL or BCL-2 alone.(17, 20, 22–24) Additionally, our study indicates delayed resistance may arise in response to dual BCL-xL/2 degrader, which may not be due to non-degradation of BCL-xL and BCL-2. This might be attributed to the capacity of cancer cells to adapt to therapeutic pressure such as compensatory activation of other pro-survival pathways. Therefore, future research should focus on identifying strategies to overcome potential resistance mechanisms, such as exploring combination therapies. In conclusion, our study provides compelling evidence for the therapeutic efficacy of co-targeting BCL-xL and BCL-2 using PROTAC degraders in SCLC. Early-phase clinical trials evaluating the safety and efficacy of navitoclax in SCLC patients have shown encouraging efficacy, underscoring the translational potential of this therapeutic approach. However, navitoclax showed severe reductions in platelet levels in patients, which hinders its clinical translation.(27) Therefore, the BCL-xL/2 dual degraders warrant clinical testing, which have high potential of enhanced efficacy and/or reduced toxicity.

We have demonstrated a potent pro-apoptotic response and laid the foundation for further clinical development of this innovative therapeutic strategy. As we move forward, a multidisciplinary approach encompassing preclinical investigations, translational research, and clinical trials will be instrumental in realizing the full potential of dual BCL-xL/2 degradation in improving outcomes for SCLC patients.

## Acknowledgments

Funding from US National Institutes of Health (NIH) grant R01 CA242003 (D.Z. & G.Z.), R01 CA241191 (D.Z. & G.Z.), and the Mike Hogg Fund (S.K.).

## Author contributions

S.K. Conceived, designed, and supervised the study, designed and performed most of the experiments, analyzed and interpreted data, and wrote and revised the manuscript; L.C. performed some of the immunoblotting experiments; J.W. assisted in xenograft experiments; P.Z. synthesized and purified DT2216 and 753b and prepared the vehicle and formulated DT2216 and 753b for the in vivo studies; G.Z. supervised the synthesis, purification, and formulation of DT2216 and 753b, and revised the manuscript; M.Z.-K. and F.J.K. provided well characterized SCLC cell lines, revised and commented on the manuscript; D.Z. Co-supervised the study, acquired funding and other resources and revised the manuscript. All authors discussed the results and commented on the manuscript.

## REFERENCES

1. Liang J, Guan X, Bao G, Yao Y, Zhong X. Molecular subtyping of small cell lung cancer. Semin Cancer Biol. 2022;86(Pt 2):450–62.

2. Megyesfalvi Z, Gay CM, Popper H, Pirker R, Ostoros G, Heeke S, et al. Clinical insights into small cell lung cancer: Tumor heterogeneity, diagnosis, therapy, and future directions. CA Cancer J Clin. 2023;73(6):620–52.

3. Rudin CM, Brambilla E, Faivre-Finn C, Sage J. Small-cell lung cancer. Nat Rev Dis Primers. 2021;7(1):3.

4. Zugazagoitia J, Paz-Ares L. Extensive-Stage Small-Cell Lung Cancer: First-Line and Second-Line Treatment Options. J Clin Oncol. 2022;40(6):671–80.

5. Mathieu L, Shah S, Pai-Scherf L, Larkins E, Vallejo J, Li X, et al. FDA Approval Summary: Atezolizumab and Durvalumab in Combination with Platinum-Based Chemotherapy in Extensive Stage Small Cell Lung Cancer. Oncologist. 2021;26(5):433–8.

6. Horn L, Mansfield AS, Szczesna A, Havel L, Krzakowski M, Hochmair MJ, et al. First-Line Atezolizumab plus Chemotherapy in Extensive-Stage Small-Cell Lung Cancer. N Engl J Med. 2018;379(23):2220–9.

7. Markham A. Lurbinectedin: First Approval. Drugs. 2020;80(13):1345–53.

8. Singh S, Jaigirdar AA, Mulkey F, Cheng J, Hamed SS, Li Y, et al. FDA Approval Summary: Lurbinectedin for the Treatment of Metastatic Small Cell Lung Cancer. Clin Cancer Res. 2021;27(9):2378–82.

9. Desai A, Smith CJ, Ashara Y, Orme JJ, Zanwar S, Potter A, et al. Real-World Outcomes With Lurbinectedin in Second-Line Setting and Beyond for Extensive Stage Small Cell Lung Cancer. Clin Lung Cancer. 2023;24(8):689–95 e1.

10. Ashkenazi A, Fairbrother WJ, Leverson JD, Souers AJ. From basic apoptosis discoveries to advanced selective BCL-2 family inhibitors. Nat Rev Drug Discov. 2017;16(4):273–84.

11. Singh R, Letai A, Sarosiek K. Regulation of apoptosis in health and disease: the balancing act of BCL-2 family proteins. Nat Rev Mol Cell Biol. 2019;20(3):175–93.

12. Lochmann TL, Floros KV, Naseri M, Powell KM, Cook W, March RJ, et al. Venetoclax Is Effective in Small-Cell Lung Cancers with High BCL-2 Expression. Clin Cancer Res. 2018;24(2):360–9.

13. Shoemaker AR, Mitten MJ, Adickes J, Ackler S, Refici M, Ferguson D, et al. Activity of the Bcl-2 family inhibitor ABT-263 in a panel of small cell lung cancer xenograft models. Clin Cancer Res. 2008;14(11):3268–77.

14. Tahir SK, Wass J, Joseph MK, Devanarayan V, Hessler P, Zhang H, et al. Identification of expression signatures predictive of sensitivity to the Bcl-2 family member inhibitor ABT-263 in small cell lung carcinoma and leukemia/lymphoma cell lines. Mol Cancer Ther. 2010;9(3):545–57.

15. Tse C, Shoemaker AR, Adickes J, Anderson MG, Chen J, Jin S, et al. ABT-263: a potent and orally bioavailable Bcl-2 family inhibitor. Cancer Res. 2008;68(9):3421–8.

16. Faber AC, Farago AF, Costa C, Dastur A, Gomez-Caraballo M, Robbins R, et al. Assessment of ABT-263 activity across a cancer cell line collection leads to a potent combination therapy for small-cell lung cancer. Proc Natl Acad Sci U S A. 2015;112(11):E1288–96.

17. Gardner EE, Connis N, Poirier JT, Cope L, Dobromilskaya I, Gallia GL, et al. Rapamycin rescues ABT-737 efficacy in small cell lung cancer. Cancer Res. 2014;74(10):2846–56.

18. Inoue-Yamauchi A, Jeng PS, Kim K, Chen HC, Han S, Ganesan YT, et al. Targeting the differential addiction to anti-apoptotic BCL-2 family for cancer therapy. Nat Commun. 2017;8:16078.

19. Li H, Wang H, Deng K, Han W, Hong B, Lin W. The ratio of Bcl-2/Bim as a predictor of cisplatin response provides a rational combination of ABT-263 with cisplatin or radiation in small cell lung cancer. Cancer Biomark. 2019;24(1):51–9.

20. Potter DS, Galvin M, Brown S, Lallo A, Hodgkinson CL, Blackhall F, et al. Inhibition of PI3K/BMX Cell Survival Pathway Sensitizes to BH3 Mimetics in SCLC. Mol Cancer Ther. 2016;15(6):1248–60.

21. Yasuda Y, Ozasa H, Kim YH, Yamazoe M, Ajimizu H, Yamamoto Funazo T, et al. MCL1 inhibition is effective against a subset of small-cell lung cancer with high MCL1 and low BCL-X(L) expression. Cell Death Dis. 2020;11(3):177.

22. Mattoo AR, FitzGerald DJ. Combination treatments with ABT-263 and an immunotoxin produce synergistic killing of ABT-263-resistant small cell lung cancer cell lines. Int J Cancer. 2013;132(4):978–87.

23. Nakajima W, Sharma K, Hicks MA, Le N, Brown R, Krystal GW, Harada H. Combination with vorinostat overcomes ABT-263 (navitoclax) resistance of small cell lung cancer. Cancer Biol Ther. 2016;17(1):27–35.

24. Wang H, Hong B, Li X, Deng K, Li H, Yan Lui VW, Lin W. JQ1 synergizes with the Bcl-2 inhibitor ABT-263 against MYCN-amplified small cell lung cancer. Oncotarget. 2017;8(49):86312–24.

25. Valko Z, Megyesfalvi Z, Schwendenwein A, Lang C, Paku S, Barany N, et al. Dual targeting of BCL-2 and MCL-1 in the presence of BAX breaks venetoclax resistance in human small cell lung cancer. Br J Cancer. 2023;128(10):1850–61.

26. Khan S, Kellish P, Connis N, Thummuri D, Wiegand J, Zhang P, et al. Co-targeting BCL-X(L) and MCL-1 with DT2216 and AZD8055 synergistically inhibit small-cell lung cancer growth without causing on-target toxicities in mice. Cell Death Discov. 2023;9(1):1.

27. Rudin CM, Hann CL, Garon EB, Ribeiro de Oliveira M, Bonomi PD, Camidge DR, et al. Phase II study of single-agent navitoclax (ABT-263) and biomarker correlates in patients with relapsed small cell lung cancer. Clin Cancer Res. 2012;18(11):3163–9.

28. Schoenwaelder SM, Jarman KE, Gardiner EE, Hua M, Qiao J, White MJ, et al. Bcl-xL-inhibitory BH3 mimetics can induce a transient thrombocytopathy that undermines the hemostatic function of platelets. Blood. 2011;118(6):1663–74.

29. Zhang H, Nimmer PM, Tahir SK, Chen J, Fryer RM, Hahn KR, et al. Bcl-2 family proteins are essential for platelet survival. Cell Death Differ. 2007;14(5):943–51.

30. Negi A, Voisin-Chiret AS. Strategies to Reduce the On-Target Platelet Toxicity of Bcl-x(L) Inhibitors: PROTACs, SNIPERs and Prodrug-Based Approaches. Chembiochem. 2022;23(12):e202100689.

31. Lakhani NJ, Rasco D, Wang H, Men L, Liang E, Fu T, et al. First-in-Human Study with Preclinical Data of BCL-2/BCL-xL Inhibitor Pelcitoclax in Locally Advanced or Metastatic Solid Tumors. Clin Cancer Res. 2024;30(3):506–21.

32. Khan S, Zhang X, Lv D, Zhang Q, He Y, Zhang P, et al. A selective BCL-X(L) PROTAC degrader achieves safe and potent antitumor activity. Nat Med. 2019;25(12):1938–47.

33. He Y, Koch R, Budamagunta V, Zhang P, Zhang X, Khan S, et al. DT2216-a Bcl-xL-specific degrader is highly active against Bcl-xL-dependent T cell lymphomas. J Hematol Oncol. 2020;13(1):95.

34. Zhang P, Zhang X, Liu X, Khan S, Zhou D, Zheng G. PROTACs are effective in addressing the platelet toxicity associated with BCL-X(L) inhibitors. Explor Target Antitumor Ther. 2020;1(4):259–72.

35. Zhang X, Thummuri D, Liu X, Hu W, Zhang P, Khan S, et al. Discovery of PROTAC BCL-X(L) degraders as potent anticancer agents with low on-target platelet toxicity. Eur J Med Chem. 2020;192:112186.

36. Lv D, Pal P, Liu X, Jia Y, Thummuri D, Zhang P, et al. Development of a BCL-xL and BCL-2 dual degrader with improved anti-leukemic activity. Nat Commun. 2021;12(1):6896.

37. Phelps RM, Johnson BE, Ihde DC, Gazdar AF, Carbone DP, McClintock PR, et al. NCI-Navy Medical Oncology Branch cell line data base. J Cell Biochem Suppl. 1996;24:32–91.

38. Iorio F, Knijnenburg TA, Vis DJ, Bignell GR, Menden MP, Schubert M, et al. A Landscape of Pharmacogenomic Interactions in Cancer. Cell. 2016;166(3):740–54.

39. Ryu Y, Hall CP, Reynolds CP, Kang MH. Caspase-dependent Mcl-1 cleavage and effect of Mcl-1 phosphorylation in ABT-737-induced apoptosis in human acute lymphoblastic leukemia cell lines. Exp Biol Med (Maywood). 2014;239(10):1390–402.

40. Wang T, Yang Z, Zhang Y, Zhang X, Wang L, Zhang S, Jia L. Caspase cleavage of Mcl-1 impairs its anti-apoptotic activity and proteasomal degradation in non-small lung cancer cells. Apoptosis. 2018;23(1):54–64.

41. Weng C, Li Y, Xu D, Shi Y, Tang H. Specific cleavage of Mcl-1 by caspase-3 in tumor necrosis factor-related apoptosis-inducing ligand (TRAIL)-induced apoptosis in Jurkat leukemia T cells. J Biol Chem. 2005;280(11):10491–500.

42. Deeks ED. Venetoclax: First Global Approval. Drugs. 2016;76(9):979–87.

43. Guerra VA, DiNardo C, Konopleva M. Venetoclax-based therapies for acute myeloid leukemia. Best Pract Res Clin Haematol. 2019;32(2):145–53.

