## Supplementary Information for "PROTAC-mediated dual degradation of BCL-xL and BCL-2 is a highly effective therapeutic strategy in small-cell lung cancer"

**Khan *et al*.**

**SUPPLEMENTARY FIGURES**

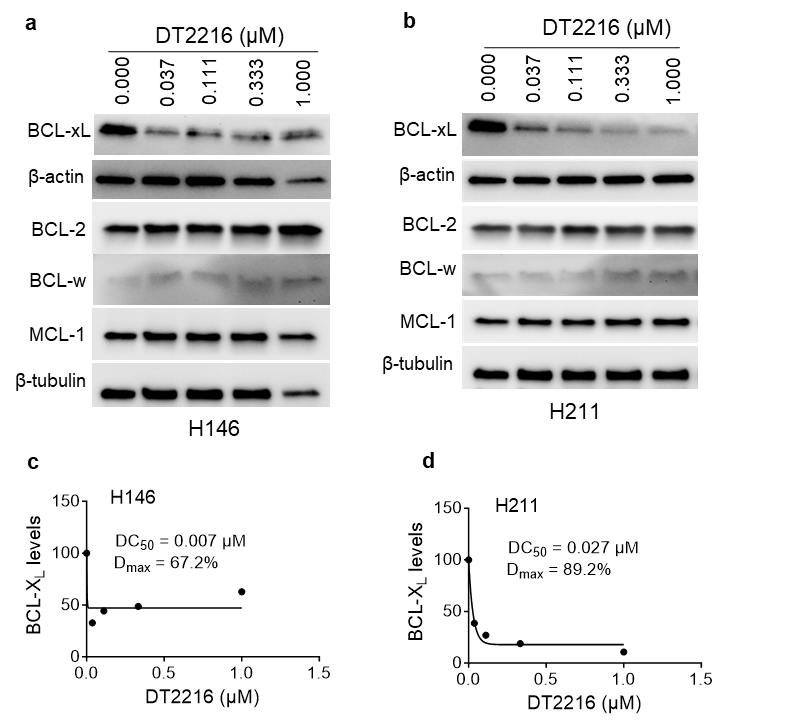

**Supplementary Fig. 1. DT2216 degrades BCL-X_L_ in SCLC cells**. **a,b,** Immunoblot analyses of BCL-X_L_ in H146 (a) and H211 (b) cells after they were treated with increasing concentrations of DT2216 for 48 h. β-actin was used as an equal loading control. **c,d**, Densitometric analysis of BCL-X_L_ along with DC_50_ and D_max_ values in H146 (c) and H211 (d) cells are shown.

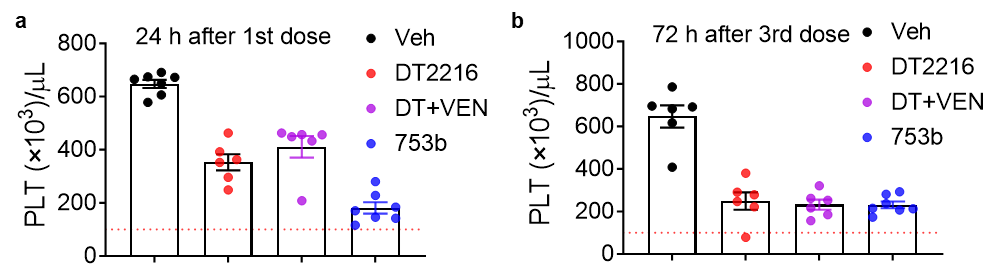

**Supplementary Fig. 2. 753b did not cause appreciable platelet toxicity in mice. a**, Enumeration of platelets in mice blood 24 h after the 1^st^ dose with vehicle, DT2216, DT2216+venetoclax, or 753b. Data are presented as mean ± SEM (n = 7, 6 and 7 mice in vehicle, DT2216 and 753b groups, respectively). **b**, Enumeration of platelets in mice blood 72 h after the 3^rd^ weekly dose with vehicle, DT2216, DT2216+venetoclax, or 753b. Data are presented as mean ± SEM (n = 6, 6 and 7 mice in vehicle, DT2216 and 753b groups, respectively).

**Supplementary Table 1. Antibodies used in immunoblotting**

| **Antibody** | **Clone** | **Antibody isotype** | **Catalog #** | **Concentration** |
| --- | --- | --- | --- | --- |
| BCL-X_L_ | _ | Rabbit IgG polyclonal | 2762 | 1:1000 |
| BCL-2 | 50E3 | Rabbit IgG monoclonal | 2870 | 1:500 |
| MCL-1 | D35A5 | Rabbit IgG monoclonal | 5453 | 1:1000 |
| BCL-w | 31H4 | Rabbit IgG monoclonal | 2724 | 1:500 |
| PARP/Cleaved PARP | 46D11 | Rabbit IgG monoclonal | 9532 | 1:1000 |
| Full-length caspase-3 | _ | Rabbit IgG polyclonal | 9662 | 1:1000 |
| cleaved caspase-3 | _ | Rabbit IgG polyclonal | 9661 | 1:1000 |
| β-tubulin | _ | Rabbit IgG polyclonal | 2146 | 1:3000 |
| β-actin | D6A8 | Rabbit IgG monoclonal | 8457 | 1:5000 |
| Secondary antibody |  | Anti-rabbit IgG, HRP | 7074 | 1:3000 |

**Footnotes:** All the antibodies were purchased from Cell Signaling Technology, Danvers, MA.
